# MitoVisualize: A resource for analysis of variants in human mitochondrial RNAs and DNA

**DOI:** 10.1101/2021.12.04.470997

**Authors:** Nicole J. Lake, Lily Zhou, Jenny Xu, Monkol Lek

**Affiliations:** Department of Genetics, Yale School of Medicine, New Haven, CT, USA; Murdoch Children’s Research Institute, Melbourne, Australia

## Abstract

We present MitoVisualize, a new tool for analysis of the human mitochondrial DNA (mtDNA). MitoVisualize enables visualization of: (1) the position and effect of variants in mitochondrial transfer RNA (tRNA) and ribosomal RNA (rRNA) secondary structures alongside curated variant annotations, (2) data across RNA structures, such as to show all positions with disease-associated variants or with post-transcriptional modifications, and (3) the position of a base, gene or region in the circular mtDNA map, such as to show the location of a large deletion. All visualizations can be easily downloaded as figures for reuse. MitoVisualize can be useful for anyone interested in exploring mtDNA variation, though is designed to facilitate mtDNA variant interpretation in particular. MitoVisualize can be accessed via https://www.mitovisualize.org/. The source code is available at https://github.com/leklab/mito_visualize/.

## INTRODUCTION

The human mitochondrial DNA (mtDNA) is a small but essential part of our genome, and encodes for proteins as well as ribosomal RNAs (rRNA) and transfer RNAs (tRNA) (Anderson *et al*., 1981). Pathogenic mtDNA variants cause a range of phenotypes collectively known as mitochondrial diseases (Gorman *et al*., 2016). Mitochondrial variants are also involved in other diseases, such as neurological disorders and cancer, and in various traits related to health (Keogh and Chinnery, 2015; Gammage and Frezza, 2019; Yonova-Doing *et al*., 2021). However the effect of most mtDNA variants on mitochondrial function, and their role in health and disease, remains unknown.

Several tools have been developed for mtDNA variant analysis, such as MITOMASTER, HmtVar and mvTool, which provide annotations in tabular format (Lott *et al*., 2013; Preste *et al*., 2019; Shen *et al*., 2018). MitImpact provides annotations for nonsynonymous variants, including a visualization in the protein structure, but does not cover RNAs (Castellana *et al*., 2015). For RNA genes the position of a variant in the secondary structure, and its effect on base pairing, are important considerations when evaluating its impact (McFarland *et al*., 2004). Analysis of data across the RNA structure, such as to show bases with pathogenic variants, is also useful (Pütz *et al*., 2007). However tools for visualizing variants and data within RNA structures had not been developed; nor had tools for generating figures labelling bases and regions within the circular mtDNA map.

To address this we developed MitoVisualize, a new tool for visualization of variants and data in RNA secondary structures, and of loci in the circular mtDNA.

## IMPLEMENTATION

Various curated data are displayed on MitoVisualize. The secondary structures and domain annotations for the 22 tRNAs are per Mamit-tRNA (Pütz *et al*., 2007), and as published for the 2 rRNA secondary structures (Amunts *et al*., 2015; Brown *et al*., 2014). The mtDNA map is NCBI Reference Sequence NC_012920.1 (Andrews *et al*., 1999). Annotations provided include (1) population frequency from gnomAD (Laricchia *et al*., 2021), MITOMAP (Lott *et al*., 2013) and HelixMTdb (Bolze *et al*., 2020) databases, (2) maximum heteroplasmy from gnomAD and HelixMTdb, (3) tRNA *in silico* predictions from MitoTIP (Sonney *et al*., 2017), PON-mt-tRNA (Niroula and Vihinen, 2016), and HmtVar (Preste *et al*., 2019), (4) disease associations, and their classification status, from MITOMAP and ClinVar (Landrum *et al*., 2018), (5) associated haplogroups from Phylotree (van Oven and Kayser, 2009), (5) conservation scores from PhyloP and PhastCons (Pollard *et al*., 2010), and (6) other data types such as bases with post-transcriptional modifications (Suzuki *et al*., 2020; Rebelo-Guiomar *et al*., 2019) or involved in formation of the tertiary structures (tRNA only, from HmtVar). Detailed information on data collection and display is in the Supplementary Table.

MitoVisualize was built using open source tools. On the server end it runs an Elasticsearch database with data populated from sources described above. Users and other applications can access this processed data directly via the GraphQL interface (https://mitovisualize.org/api). The web front-end was implemented in JavaScript using React libraries. Lastly, the RNA and mtDNA visualizations were custom designed in SVG format.

## USING MITOVISUALIZE

A search bar for querying mitochondrial RNA variants and genes is located on the landing page, as well as in the navigation bar accessible from every page. The About page summarizes key functionality and data used to build MitoVisualize, and ‘?’ icons throughout the site provide extra information about the various annotations. There are three key functions in MitoVisualize: (1) using the variant tool to analyze a RNA variant, (2) using the gene tool to visualize data across RNA structures, and (3) using the mtDNA tool to annotate a base, gene or region in the circular mtDNA.

The variant tool can be accessed by typing the RNA variant of interest into the search bar, or by selecting ‘Variant’ in the dropdown menu for tRNA and rRNA in the navigation bar. MitoVisualize currently only accepts single nucleotide variants (input format e.g. m.3243A>G). The position of the variant within the RNA secondary structure is shown in red text and yellow highlight; any change to a base pair type is also displayed (Figure 1A). For rRNAs the region with the variant is shown zoomed-in, while its location in the larger structure is displayed in the corner inset. Various annotations are provided alongside the RNA structure visualization to help the user interpret the impact of the variant (see Supplementary Table). All annotations are hyperlinked for easy navigation to the data source.

**Figure 1:**
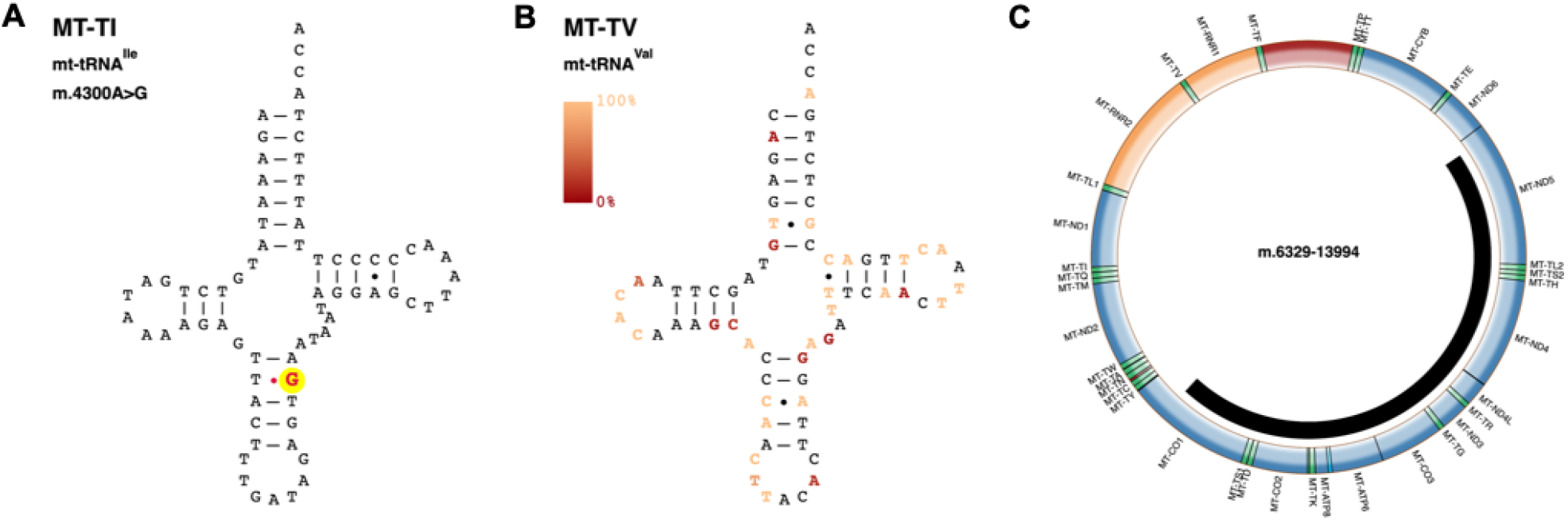
Example visualizations in MitoVisualize. **(A)** Using the variant tool to visualize the position and effect of m.4300A>G. **(B)** Using the gene tool to visualize maximum heteroplasmy in gnomAD across *MT-TV*. **(C)** Using the mtDNA tool to visualize a deletion spanning m.6329-13994.

The gene tool can be accessed by typing the RNA gene of interest into the search bar, or by selecting ‘Gene’ in the dropdown menu for tRNA and rRNA in the navigation bar. A user can select a data type of interest for visualization across the RNA structure; note the mtDNA coordinate of a base can be revealed by hovering over it. Quantitative data types (i.e. maximum heteroplasmy of base variants in population databases, Figure 1B) are visualized as a color gradient, while qualitative data (i.e. bases with disease-associated variants) are shown in red.

The mtDNA tool is accessible via the navigation bar and displays the circular mtDNA molecule. Protein-coding genes are colored blue, rRNA genes yellow, tRNA genes in green, and non-coding regions in red. A user can label the position of a gene, base/variant, or region within the mtDNA (Figure 1C). For regions, overlapping genes are also listed.

## SUMMARY

MitoVisualize is a free, user-friendly tool for visualizing the position and effect of variants in human mitochondrial tRNA and rRNA secondary structures, and in the circular mtDNA map. It also enables visualization of data across the RNA secondary structures. All visualizations can be downloaded as png figures for reuse. By including the variant or gene name in the URL, we enable easy linking to MitoVisualize from other resources; users can also access data directly via API. Version and date of access for all data sources are listed on the About page of MitoVisualize, and will be updated periodically.

## ACKNOWLEDGEMENTS

We gratefully acknowledge the contribution of Tianxiao (Lisa) Wang who helped create the RNA SVG files.

## FUNDING

This work was supported by the National Health and Medical Research Council [1159456 to N.J.L.].

## Conflict of interest

None declared.

## Notes

https://www.mitovisualize.org/

### Competing Interest Statement

The authors have declared no competing interest.

