## Supplementary Material for "MitoVisualize: A resource for analysis of variants in human mitochondrial RNAs and DNA"

**Supplementary Table:** The different data displayed on MitoVisualize, and notes on their collection and display, is described. Version information and date of access for all data is also listed on the About page of the website - https://www.mitovisualize.org/about-page.

| **Data type** | | **Data sources** | **Data collection notes** | **Display notes** | **References** |
|---|---|---|---|---|---|
| Structural | mtDNA molecule | National Center for Biotechnology Information (NCBI) | NCBI Reference NC_012920.1 (revised Cambridge reference sequence, rCRS) | Displayed in counter-clockwise orientation, the traditional map representation | (Andrews *et al.*, 1999) |
| | RNA secondary structures | Mamit-tRNA (tRNA) | The structures for all 22 tRNAs and 2 rRNAs were manually drawn as Scalable Vector Graphic (SVG) images | Watson-Crick (WC) base pairs in the double-stranded stems are denoted by '-', while non-WC pairs are represented by '•' | (Pütz *et al.*, 2007) |
| | | Amunts et al, Brown et al (rRNA) | | | (Amunts *et al.*, 2015; Brown *et al.*, 2014) |
| | Domain annotations | Mamit-tRNA | Per the nucleotide numbering table at mamit-trna.u-strasbg.fr/human.asp, manually transcribed | tRNA only | (Pütz *et al.*, 2007) |
| Variant | Population frequency (and allele count) | gnomAD | Version 3.1 | Shown separately for homoplasmic and heteroplasmic variants | (Laricchia *et al.*, 2021) |
| | | HelixMTdb | Version dated 03/27/2020 | | (Bolze *et al.*, 2020) |
| | | MITOMAP | Date of access 08/31/2021 | - | (Lott *et al.*, 2013) |
| | Maximum heteroplasmy | gnomAD | Version 3.1 | - | (Laricchia *et al.*, 2021) |
| | | HelixMTdb | Version dated 03/27/2020 | Not reported for homoplasmic variants, which was assigned to be 100% | (Bolze *et al.*, 2020) |

| | | | | | |
|---|---|---|---|---|---|
| | In silico predictions | MitoTip | Retrieved from MITOMAP | tRNA only | (Sonney *et al.*, 2017) |
| | | PON-mt-tRNA | Retrieved from structure.bmc.lu.se/PON-mt-tRNA/ | | (Niroula and Vihinen, 2016) |
| | | HmtVar | Retrieved from HmtDB via API | | (Preste *et al.*, 2019) |
| | Disease association | MITOMAP | Date of access 08/31/2021 | For MITOMAP the ‘status’ is shown, for ClinVar the ‘classification’ is shown | (Lott *et al.*, 2013) |
| | | ClinVar | Date of access 08/31/2021 | | (Landrum *et al.*, 2018) |
| | Associated haplogroups | Phylotree | Extracted from PhyloTree Build 17 using custom scripts | - | (van Oven and Kayser, 2009) |
| Base | Conservation | PhyloP | Conservation scores derived from 100 vertebrate genomes, retrieved via the UCSC genome browser | - | (Pollard *et al.*, 2010) |
| | | PhastCons | | | |
| | Post-transcriptionally modified bases | Suzuki et al (tRNA) | tRNA position numbers of modifications were manually converted to the corresponding mtDNA coordinates | - | (Suzuki *et al.*, 2020) |
| | | Rebelo-Guiomar et al (rRNA) | Manually transcribed the corresponding mtDNA coordinates | - | (Rebelo-Guiomar *et al.*, 2019) |
| | 3D interactions (folding) | HmtVar | Retrieved from HmtDB via API | tRNA only | (Preste *et al.*, 2019) |
